# Mice harboring a SCA28 patient mutation in *AFG3L2* develop late-onset ataxia associated with enhanced mitochondrial proteotoxicity

**DOI:** 10.1101/300202

**Authors:** Cecilia Mancini, Eriola Hoxha, Luisa Iommarini, Alessandro Brussino, Uwe Richter, Francesca Montarolo, Claudia Cagnoli, Roberta Parolisi, Diana Iulia Gondor Morosini, Valentina Nicolò, Francesca Maltecca, Luisa Muratori, Giulia Ronchi, Stefano Geuna, Francesca Arnaboldi, Elena Donetti, Elisa Giorgio, Simona Cavalieri, Eleonora Di Gregorio, Elisa Pozzi, Marta Ferrero, Evelise Riberi, Giorgio Casari, Fiorella Altruda, Emilia Turco, Giuseppe Gasparre, Brendan J. Battersby, Anna Maria Porcelli, Enza Ferrero, Alfredo Brusco, Filippo Tempia

## Abstract

Spinocerebellar ataxia 28 is an autosomal dominant neurodegenerative disorder caused by missense mutations affecting the proteolytic domain of AFG3L2, a major component of the mitochondrial *m*-AAA protease. However, little is known of the underlying pathogenetic mechanisms or how to treat patients with SCA28. Currently available *Afg3l2* mutant mice harbour deletions that lead to severe, early-onset neurological phenotypes that do not faithfully reproduce the late-onset and slowly progressing SCA28 phenotype. Here we describe production and detailed analysis of a new knock-in murine model harbouring an *Afg3l2* allele carrying the p.Met665Arg patient-derived mutation. Heterozygous mutant mice developed normally but signs of ataxia were detectable by beam test at 18 months. Cerebellar pathology was negative; electrophysiological analysis showed increased spontaneous firing in Purkinje cells from heterozygous mutants with respect to wild-type controls, although not statistically significant. As homozygous mutants died perinatally with evidence of cardiac atrophy, for each genotype we generated mouse embryonic fibroblasts (MEFs) to investigate mitochondrial function. MEFs from mutant mice showed altered mitochondrial bioenergetics, with decreased basal oxygen consumption rate, ATP synthesis and mitochondrial membrane potential. Mitochondrial network formation and morphology was also altered, in line with greatly reduced expression of Opa1 fusogenic protein L-isoforms. The mitochondrial alterations observed in MEFs were also detected in cerebella of 18-month-old heterozygous mutants, suggesting they may be a hallmark of disease. Pharmacological inhibition of *de novo* mitochondrial protein translation with chloramphenicol caused reversal of mitochondrial morphology in homozygous mutant MEFs, supporting the relevance of mitochondrial proteotoxicity for SCA28 pathogenesis and therapy development.

## INTRODUCTION

The hereditary spinocerebellar ataxias (SCAs) are a group of over 40 neurodegenerative disorders characterized by autosomal dominant inheritance (Durr 2010). Although each form of SCA has its own distinct causative gene, the pathogenetic pathways converge on cerebellar and spinal degeneration leading to an array of slowly progressive neurological deficits (Durr 2010; Nibbeling et al. 2017; Smeets and Verbeek 2014). We first described SCA type 28 (SCA28) over a decade ago in a four-generation Italian family with ataxia (Cagnoli et al. 2006; Di Bella et al. 2010; Mariotti et al. 2008), and later identified mutations in *AFG3L2* (AFG3 ATPase Family Member 3-Like 2) as the cause of disease (Di Bella et al. 2010). Clinically, the mean age of onset was 19.5 years with signs of altered balance and gait. Limb ataxia, dysarthria and eye movement abnormalities were also observed. The disease course is very slow, and patients remain ambulant into their late sixties. The principal finding by brain MRI is cerebellar atrophy, that usually appears as first hallmark of the disease.

SCA28 is a rare disease, representing ∼1.5% of all autosomal dominant cerebellar ataxias (whose prevalence is 1-9/100,000). SCA28 has no treatment options, also because we lack a mechanistic understanding of disease pathogenesis. SCA28 is unique in being currently the only form of dominant ataxia caused by dysfunction of a mitochondrial-dwelling protein (http://neuromuscular.wustl.edu/ataxia/domatax.html). The causative gene *AFG3L2* is part of the nuclear genome and encodes a protein subunit that forms the *m*-AAA protease (matrix-ATPase associated with diverse cellular activities), a multimeric complex bound to the inner mitochondrial membrane (Koppen et al. 2007). The *m*-AAA protease is part of a complex network of evolutionarily conserved proteases that represent the most important inner defence system of mitochondrial integrity (Koppen and Langer 2007; Patron et al. 2018), which is maintained also by other mechanisms such as regeneration or culling of compromised mitochondria by mitochondrial dynamics (fusion and fission) and removal of damaged organelles by mitophagy (Liesa et al. 2009). The *m-*AAA protease is a crucial component of the mitochondrial protein quality control system, exerting a chaperone-like activity during biogenesis of oxidative phosphorylation (OXPHOS) respiratory chain complexes. The *m*-AAA also participates in mitochondrial protein processing and maturation (Arlt et al. 1998; Atorino et al. 2003; Gerdes et al. 2012; Koppen and Langer 2007; Nolden et al. 2005).

In humans, AFG3L2 is a major constituent of the *m*-AAA protease, being capable of self-aggregation to form the homo-hexameric form of *m*-AAA. AFG3L2 can also form hetero-hexamers in partnership with its paralogue SPG7 (alias paraplegin) (Koppen et al. 2007). To become active, both AFG3L2 and SPG7 themselves need to be proteolytically processed. However, while AFG3L2 is capable of self-activation, SPG7 requires AFG3L2 for maturation and activation via tyrosine phosphorylation, and does not self-assemble (Almontashiri et al. 2014; Koppen et al. 2009). Mutations in SPG7 are associated with the genetic disorder hereditary spastic paraplegia type 7 (Casari et al. 1998).

Structurally, the *m*-AAA protease is essentially an ATP-fueled proteolytic machine shaped like a double-doughnut: the juxtamembrane doughnut represents the ATPase domain while the proteolytic domain protrudes into the mitochondrial matrix (Koppen & Langer 2007). Functionally, using as model the bacterial homologue FtsH, we can imagine substrates being pulled through the hexamer ring of the AAA domain into the proteolytic cleft of the protease ring, with release of degradation peptides (Dalbey et al. 2012). As yet, the identity of the human *m*-AAA substrates is unknown, but yeast *m-*AAA substrates include cytochrome *c* peroxidase (Suppanz et al. 2009) and MRPL32, a mitoribosomal protein required for ribosomal assembly and protein synthesis (Almajan et al. 2012; Nolden et al. 2005).

We dwell on the complexities of the *m*-AAA structure, function and composition because it emerges that the majority of SCA28 patient mutations induce missense changes in AFG3L2 residues that participate in the formation of the proteolytic cleft. These missense mutations are mostly clustered in *AFG3L2* exons 15 and 16 that form the protease domain: ^654^Thr→Ile; ^666^Met→Val/Arg/Thr; ^671^Gly→Arg/Glu; ^674^Ser→Leu; ^689^Tyr→Asn/His; ^691^Glu→Lys; ^694^Ala→Glu; ^700^Glu→Lys and ^702^Arg→Gln (Almajan et al. 2012; Cagnoli et al. 2010; Di Bella et al. 2010; Edener et al. 2010; Lobbe et al. 2014; Szpisjak et al. 2017; Zuhlke et al. 2015). Confirmation of the importance of modifications in the proteolytic cleft for disease pathogenesis comes from the ^432^Asn→Thr patient mutation: although encoded by exon 10, thus outside of the mutational hotspot, it modifies a cleft-facing residue from the ATPase aspect. Two pathogenic frameshift mutations have also been described in SCA28: deletion of exons 14-16 and the p.Thr654Asnfs*15, again affecting the protease domain (Musova et al. 2014; Smets et al. 2014).

In addition to these observations on the nature and location of the patients’ mutations, we note that the Exome Aggregation Consortium (ExAC) database (Lek et al. 2016) describes many human *AFG3L2* variants (synonymous, missense, frameshift, gain of stop codon and altered splice sites) that are distributed throughout all the coding region. However, the loss-of-function *AFG3L2* variants, whose frequency in the human population analysed is ∼1:2000 (*i.e.,* far higher than the incidence of SCA28) are not deleterious, and ExAC gives a probability of LoF intolerance (pLI) of 0.01, indicating that AFG3L2 is extremely tolerant to loss-of-function changes, although it must be remembered that due to the late-onset of the phenotype, it is possible that some of the mutations have not had time to become manifest. It is evident that what is not tolerated is a select group of heterozygous missense mutations that do not appear to affect AFG3L2 protein levels but instead lead to incorporation of malfunctioning AFG3L2 subunits into the *m*-AAA complexes, which in turn become dysfunctional. AFG3L2 is highly expressed in cerebellum and other neural tissues, and our working hypothesis for SCA28 pathogenesis is that mitochondrial dysfunction in these tissues leads to proteotoxic stress through the slow accumulation of toxic misfolded substrates, a process already described in other neurodegenerative diseases such as Alzheimer’s and Parkinson’s diseases (Sorrentino et al. 2017). The ensuing mitochondriopathy eventually translates into neurodegeneration and neurological manifestations.

A mouse model of SCA28 would be an ideal approach to test our mitotoxicity hypothesis. Two murine mutations affecting *Afg3l2* have already been described: one is the *paralysé* mouse (*Afg3l2^par/par^*), a spontaneous mutant, and the second is the *Afg3l2^Emv66/Emv66^* mouse, in which Afg3l2 protein is absent because of a murine leukemia proviral insertion in intron 14 (Maltecca et al. 2008). Both mutants are smaller than normal littermates at one week, show a progressive loss of limb motor function by two weeks, and are completely paralyzed and usually die by 3 weeks. Mice present severe defects in neuronal development, with axonal development failure, aberrant mitochondria, Schwann cell invagination, and impaired maturation of Purkinje cell arborization (Maltecca et al. 2008). Compound heterozygotes (*Afg3l2^par/Emv66^*) are phenotypically indistinguishable from their respective homozygotes, suggesting that both mutations act by loss of a protective function (Maltecca et al. 2008). The models have proven valuable in deciphering the functional role of Afg3l2 in axonal development, where it is required for correct assembly of respiratory chain complexes (Maltecca et al. 2008). In contrast, heterozygous *Afg3l2^Emv66/+^* mice have a milder phenotype, with the first signs of loss of balance appearing at four months and worsening with age, suggesting a dose-effect of the mutation (Maltecca et al. 2009). Additional informative models are the *Spg^7-/-^Afg3l2^Emv66/+^* digenic mutated mice obtained by crossing the *Afg3l2^Emv66/+^* strain with *SPG7* knock-out mice (Martinelli et al. 2009). These mice have a severe phenotype, characterized by early-onset loss of balance, tremor, and ataxia with altered gait coordination. Lastly, there is the *Afg3l2^PC-KO^* mouse, which limits *Afg3l2* inactivation to Purkinje cells. Again, the phenotype is severe with unsteady gait at six weeks, caused by dramatic loss of Purkinje cells (Almajan et al. 2012). All these models serve to highlight the crucial role of *Afg3l2* in maintaining normal cerebellar function, and that loss of Afg3l2 leads to disease with severe ataxic phenotypes. However, these models do not resolve the question of how the AFG3L2 missense mutations affecting the protease domain lead to SCA28.

With these considerations in mind, we hoped to generate a more authentic mouse model of disease by taking advantage of the fact that human and murine AFG3L2 proteins are highly conserved. This paved the way for knock-in generation by modifying one *Afg3l2* allele through insertion of a SCA28 patient mutation. A potential drawback of this model is that rodent genomes harbour *Afg3l1*, a paralogue of *Afg3l2* and *Spg7* that is pseudogenized in human (Kremmidiotis et al. 2001). However, *Afg3l1* shows very low expression in brain and cerebellum (Sacco et al. 2010) and does not rescue the phenotype of either *Afg3l2* or *Spg7* knock-out mice (Wang et al. 2016). Thus, we proceeded with our model, selecting the *AFG3L2* missense mutation c.1994T>G:p.Met666Arg (p.Met665Arg in mouse), reported in a paediatric case of SCA28 (Cagnoli et al. 2010). Here we describe: (i) the generation and motor behavior of the *Afg3l2^M665R^* knock-in mouse; (ii) a detailed investigation of the effects of the mutation on neural tissues and their ultrastructure; (iii) a validation of the mitochondrial proteotoxicity model and (iv) that the antibiotic chloramphenicol can revert the mitotoxic phenotype, opening a potential new avenue of treatment.

## RESULTS

### Generation of *Afg3l2^M665R/+^* knock-in mice

A targeting vector for insertion of the c.1994T>G (p.Met665Arg) missense mutation in *Afg3l2* was prepared by recombineering (Copeland et al. 2001) (Fig. 1A). Briefly, the construct contained homologous DNA arms (C57BL/6J mouse strain), a neo^*r*^ cassette and a diagnostic *Nco*I restriction site for Southern blotting. Following vector transfection into 129 embryonic stem (ES) cells, clones with one mutated allele and a single copy of neo^*r*^ were identified (Fig. 1B-C), and transferred to C57BL/6 blastocysts for germline transmission. *Afg3l2^M665R/+^* knock-in (KI) mice were born at the expected Mendelian ratios, and were normal in terms of appearance, development and fertility. Expression of both KI and wild-type (WT) alleles was verified by RT-PCR and western blotting in different tissues, including cerebellum (Fig. 1D-F) and brain (*data not shown*).

**Figure 1.**
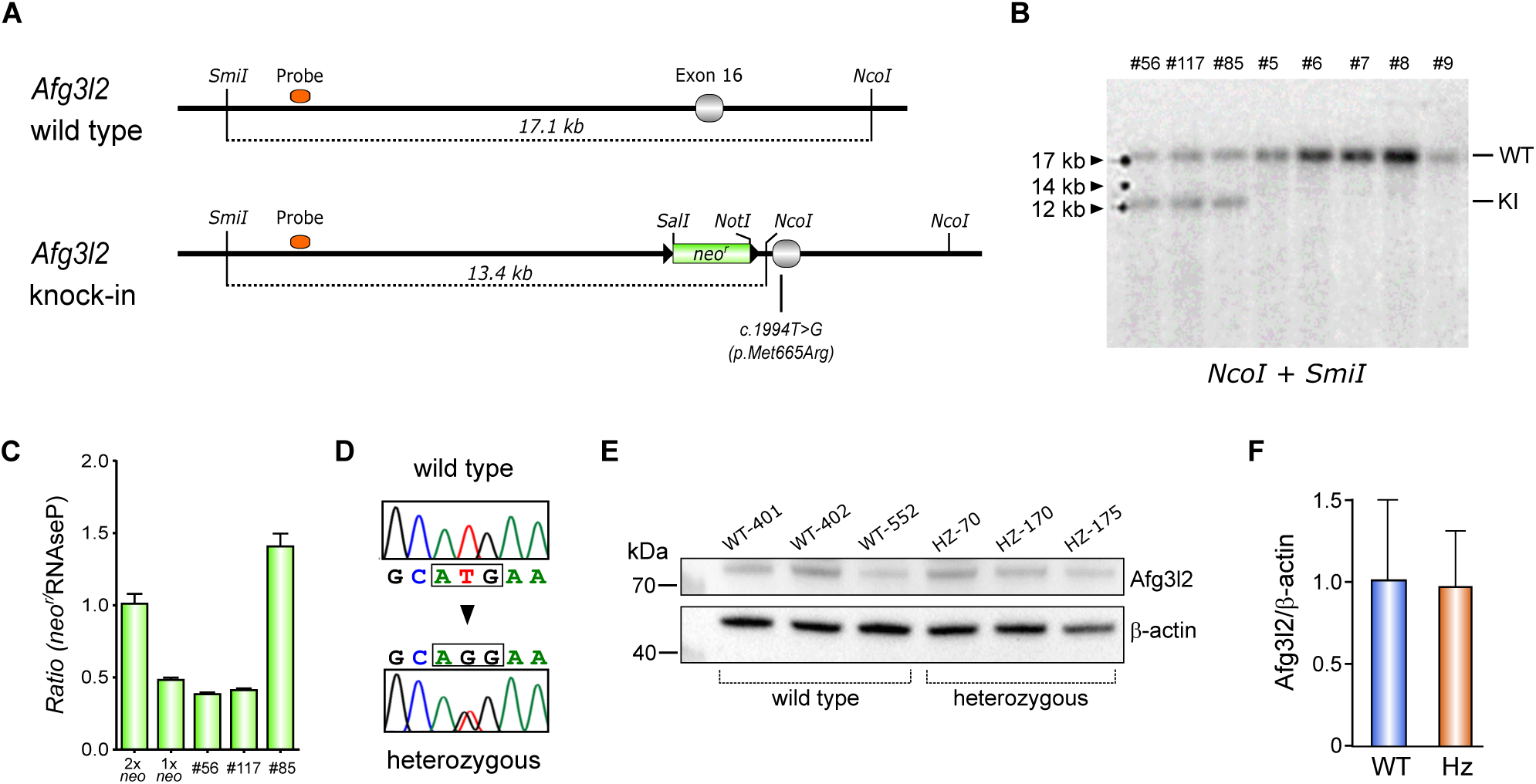
Production and molecular characterization of *Afg3l2* knock-in mice. (A). Schematic diagram of wild-type *Afg3l2* (WT) and the *Afg3l2^M665R^* (KI) allele after gene targeting by recombineering. The Southern blot probe (orange rectangle) discriminates between WT and KI alleles after *NcoI/SmiI* digestion (17.1 kb and 13.4 kb, respectively). (B). Representative Southern blot of DNA extracted from targeted 129 ES cells: DNAs #56, #117 and #85 show bands indicating the presence of both WT and KI alleles. (C). Real-time PCR quantification of *neo^r^* cassette copy number in KI-positive DNAs. DNA #56 and #117 are single copy for *neo^r^*, indicating one vector integration event; DNA #85 shows multiple copies of *neo^r^* and was discarded. The histogram represents mean ± SD of three independent experiments. (D). RT-PCR and Sanger sequencing of cerebellar *Afg3l2* cDNA from WT and heterozygous (Hz) mutant mice. (E). Detection of Afg3l2 proteins by western blot in cerebellar extracts from WT and Hz mice. (F). Histogram shows protein expression ratio between Afg3l2 and and β-actin, and represents mean ± SD of three independent experiments.

When heterozygous mutant mice were intercrossed, the expected Mendelian ratio at birth was skewed towards heterozygous and wild-type genotypes (*Afg3l2^+/+^* 39/121, *Afg3l2^M665R/+^* 75/121, *Afg3l2^M665R/M665R^* 7/121; Chi-square test, *p* = 0.0003). Most homozygous mutant mice died before or within hours of birth; rare viable mice were cyanotic. We obtained embryonic day 16.5 (E16.5) embryos for histological analyses (Supp. Fig. S1A); in *Afg3l2^M665R/M665R^* embryos, the principal finding was diffuse cardiac atrophy while cerebellar histology appeared normal (Supp. Fig. S1B-E). The cardiac phenotype and perinatal lethality of the *Afg3l2^M665R/M665R^* mouse contrasts with the *Afg3l2^Emv66/Emv66^* mouse, which dies at around 2-3 weeks after birth with a neurological (tetraparetic) and cachectic phenotype (Maltecca et al. 2008), an indication that mutations that affect AFG3L2 quality (*Afg3l2^M665R/M665R^*) and quantity (*Afg3l2^Emv66/Emv66^*) have different phenotypic effects. We return to the main focus, our hypothetical SCA28 model mouse, and leave further investigations into the cardiac phenotype of the homozygous mutant to future studies.

### Late-onset motor incoordination in *Afg3l2^M665R/+^* mice

Although there was no outer evidence of disease, we started analysing motor behavior in *Afg3l2^M665R/+^* mice and wild-type littermates at 4 months so as not to miss the onset of motor alterations. We took as a guide the fact that, at this age, *Afg3l2^Emv66/+^* mice show signs of motor incoordination. From the age of 4 months onwards, motor testing was repeated every two months. The test battery included: (i) the elevated beam test; (ii) footprint analysis and (iii) the accelerating rotarod.

When first examined (4 months), the tests showed no significant differences between wild-type and heterozygous mutants. However, at 18 months, *Afg3l2^M665R/+^* mice started to show signs of imbalance on the beam test, with significantly greater number of fore paw and hind paw slips compared to wild-type littermates. Similar results were obtained when tests were repeated on consecutive days (Fig. 2A). In contrast, no differences were observed between groups of mice in footprint analysis (Fig. 2B), in the accelerating rotarod (Fig. 2C) nor in the day 10 rotarod test (Fig. 2C), which measured motor memory retention. Since optic atrophy has been associated with *AFG3L2* mutations (Charif et al. 2015; Colavito et al. 2017), eye movement was assessed by measuring the optokinetic response in *Afg3l2^M665R/+^* and wild-type mice at 18 months. No significant differences in head tracking performances were observed between genotypes: percent still time with respect to control still time was 81.4 ± 4.41% in wild-type mice, and 90.9 ± 2.10% in heterozygous mutants. The test battery was repeated at 20, 22 and 24 months of age, with no evidence of increasing severity of the phenotype (*data not shown*). Thus, the heterozygous p.Met665Arg in Afg3l2 appears to be a mild mutation leading to adult-onset ataxia in mice that well recapitulates the SCA28 patient phenotype.

**Figure 2.**
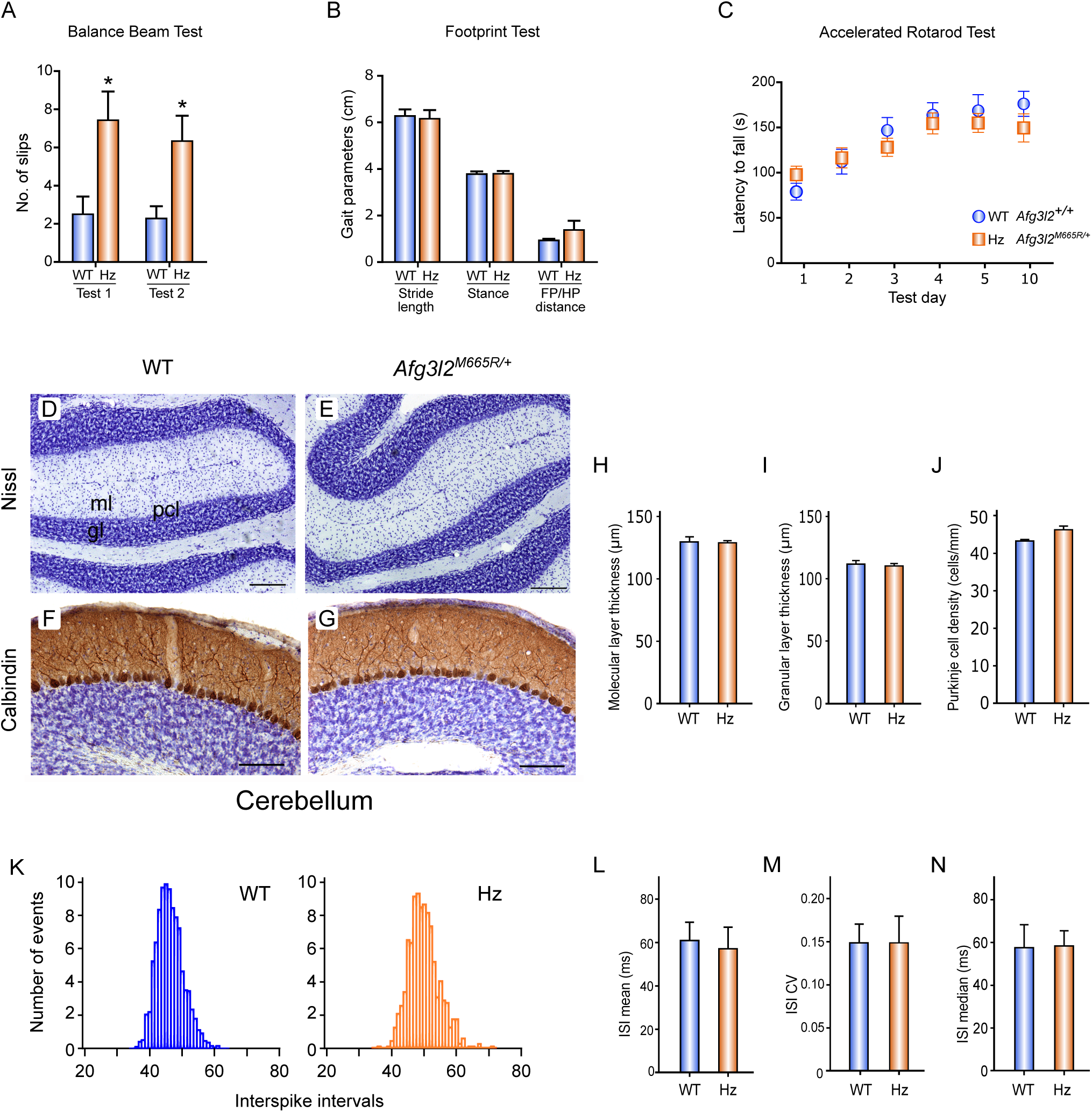
Motor test battery performance, cerebellar morphology and Purkinje cell electrophysiology of 18-month-old *Afg3l2^M665R/+^* and wild-type mice. (A). Graph shows the number of slips made on the beam test by wild-type (WT, *Afg3l2^+/+^*) and heterozygous (Hz, *Afg3l2^M665R/+^*) mice on two different test days (* is *p* < 0.05, Student’s t-test). (B). Examples of footprint analysis to evaluate gait in WT and Hz mice, showing stride length, stance and front paw/hind paw distance. Differences were not statistically significant (Student’s t-test). (C). Results of accelerated rotarod tests carried out on five consecutive test days, and on day 10. Time to fall is shown; WT and Hz mice showed similar results (repeated measures two-way ANOVA). Values are mean ± S.E.M. (D, E). Nissl-stained sagittal sections of cerebella from 18-month-old wild type (WT) and heterozygous *Afg3l2^M665R/+^* mice. Molecular layer (ml), granular layer (gl) and Purkinje cell layer (pcl) are indicated. (F, G). Calbindin-stained sagittal cerebellar sections from 18-month-old wild type and heterozygous mice. Scale bars 100 µm. (H-J). Histograms show the average thickness of the molecular (H) and granular layers (I), and Purkinje cell density (J) of the cerebellar cortex in WT and Hz mouse, showing no significant differences between genotypes (*p* > 0.05, Student’s t-test). Results are reported as mean ± SEM. Scale bars 100 µm. (K). Representative histogram showing the distribution of the interspike intervals (ISIs) in Purkinje cells from WT and Hz mice. (L-N). Histograms represent the mean ISI (L), the ISI coefficient of variation (M) and the ISI median (N) of WT (n = 24) and Hz (n = 29) Purkinje cells. No significant differences were observed between genotypes (*p* > 0.05, unpaired Student’s t-test).

### Ataxic *Afg3l2^M665R/+^* mice show normal cerebellar structure

Cerebellar degeneration is central to ataxic neuropathology and usually precedes the onset of motor dysfunction. Therefore, we began examining cerebellar structure in heterozygous mutant mice from 6 months onwards, with no pathological finding. Even in 18-month-old ataxic heterozygotes, Nissl-stained cerebellar sections showed regular lamination and foliation as in control mice (Fig. 2D, E). Molecular and granular layer thickness was quantified in all cerebellar lobules, using a previously described method (Carulli et al. 2002), with no evident differences between genotypes (Fig. 2H, I).

Since AFG3L2 is required for cerebellar Purkinje cell survival (Almajan et al. 2012), Purkinje cell numbers were estimated by calbindin staining in cerebellar sagittal sections (Fig. 2F-G). The average number of Purkinje cells was similar in *Afg3l2^M665R/+^* (46.2 ± 0.94 cells/mm; n = 3) and wild-type mice (43.2 ± 0.20 cells/mm; n = 3) (Fig. 2J). Both groups also showed similar cell numbers in the deep cerebellar nuclei (*data not shown*).

### Ataxic *Afg3l2^M665R/+^* mice show normal spontaneous firing activity in Purkinje cells

We next investigated *Afg3l2^M665R/+^* Purkinje cell function by measuring the generation of spontaneous action potentials. Individual Purkinje cells in cerebellar slices were analysed by cell-attached patch clamp recordings. The mean interspike interval (ISI) was not significantly different in Purkinje cells from 18-month-old wild-type and heterozygous mutant mice (Fig. 2K-L). The regularity of Purkinje cell firing was also conserved, as shown by the ISI coefficient of variation (Fig. 2M). The median of the interspike intervals was also similar for both genotypes (Fig. 2N). However, we did note an increase in the number of firing cells in *Afg3l2^M665R/+^* mice, but this was not statistically significant - WT Purkinje cells: 47 analysed/21 silent/26 (55%) firing; Hz Purkinje cells: 36 analysed/9 silent/27 (75%) firing.

### Normal neural ultrastructure but alterations in sciatic nerve mitochondria in *Afg3l2^M665R/+^* mice

Previously described *Afg3l2* mutant mouse models show altered mitochondrial morphology in cells of the nervous system (Maltecca et al. 2008; Maltecca et al. 2009). Therefore, we carried out detailed ultrastructural analyses of CNS- and PNS-derived tissues.

#### Cerebellum

In 18-month-old wild-type and heterozygous mutant mice, the cerebellar cortex showed the typical molecular, Purkinje and granular layer organization, with clear boundaries between layers. Purkinje cells were aligned in monolayers, with cell bodies near the granular layer, a large nucleus within the piriformis-shaped soma (Fig. 3A-B), and dendritic arborisations that crossed the molecular layer. No cell body shrinkage, enlargement of the ER *cisternae* nor fragmentation of the Golgi apparatus was observed. Mitochondria were normal with evident *cristae* and no signs of swelling (Fig. 3C-D). The granular layer was packed with small round cells (Fig. 3E-F).

**Figure 3.**
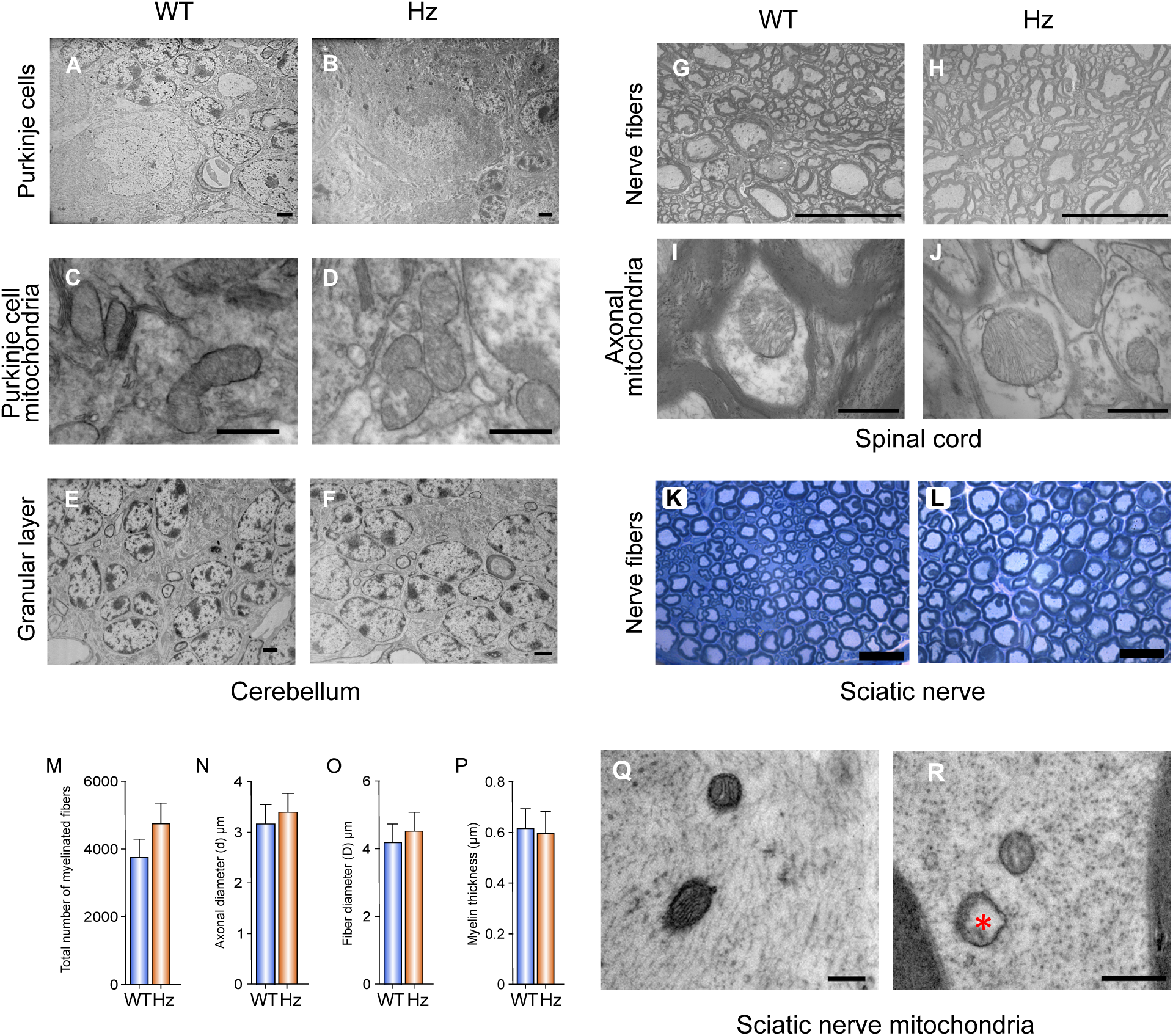
Ultrastructural analyses of cerebellum, spinal cord and sciatic nerve in wild-type and *Afg3l2^M665R/+^* mice. (A, B). Photomicrographs show ultrastructure of Purkinje cells and the inner granular layer from cerebella isolated from 18-month-old wild-type and heterozygous mutant mice. Scale bar 2 µm; (C, D). Purkinje cell mitochondria. Scale bar 500 nm; (E, F). Cells of the granular layer. Scale bar 2 µm. (G, H). Myelinated nerve fibers in the spinal cord white matter; Scale bar 20 µm; (I-J) High magnification of axonal mitochondria. Scale bar 500 nm. (K, L). Resin-embedded, 2.5 µm semithin sections of sciatic nerves derived from wild-type and *Afg3l2^M665R/+^* mice. Toluidine blue stain. Scale bar 20 µm. (M-P). Histograms of sciatic nerve stereological analysis showing comparative analyses of total numbers of myelinated fibers (M), axon diameter (N), fiber diameter (O) and myelin thickness (P). No significant differences between genotypes were observed (Student’s’ t-test). (Q, R). Representative photomicrographs of sciatic nerve obtained by transmission electron microscopy: swollen mitochondria without *cristae* are detectable in *Afg3l2^M665R/+^* mice (red asterisk). Scale bar 200 nm (Q) and 500 nm (R).

#### Spinal cord

Spinal cord white matter was similar in wild-type and *Afg3l2^M665R/+^* mice: myelinated nerve fibers of small, medium and large diameter were ubiquitous, and myelinated sheaths showed uniform thickness (Fig. 3G-H). Axonal mitochondria always displayed conserved *cristae*, homogeneous matrix, and no signs of swelling (Fig. 3I-J).

#### Sciatic nerve

Sciatic nerves were isolated and examined by high-resolution light microscopy. Nerve fibers showed similar organization and morphology, and similar size in axon diameter, fiber diameter and myelin sheaths (Fig. 3K, L). Sciatic nerve fiber morphology by stereological analysis of myelinated fibers showed no significant differences between groups in the total number of myelinated fibers, axon diameter, fiber diameter and myelin thickness (Fig. 3M-P). Increased numbers of sciatic nerve mitochondria from *Afg3l2^M665R/+^* mice showed altered *cristae* morphology and signs of swelling compared to wild-type, by stereological analysis: altered mitochondria in wild-type: 6.39 ± 1.97%; altered mitochondria in *Afg3l2^M665R/+^*: 43.1 ± 4.98%. (*p* < 0.0001, Chi-squared test) (Fig. 3Q, R). However, there were no discernible differences in peripheral nerve conduction of action potentials between groups: wild-type (n=7) 32.7 ± 1.2 m/s; *Afg3l2^M665R/+^* (n=6) 32.9 ± 1.7 m/s.

### Mitochondrial bioenergetics are altered in *Afg3l2^M665R/+^* mouse embryonic fibroblasts

To analyse the impact of the p.Met665Arg mutation on mitochondrial function, we generated mouse embryonic fibroblasts (MEFs) from E13.5 embryos as these were available for the wild-type and mutant (*Afg3l2^M665R/+^* and *Afg3l2^M665R/M665R^*) genotypes. MEFs were then used to analyse mitochondrial dynamics, bioenergetics, and oxidative phosphorylation (OXPHOS). Compared to wild-type MEFs, *Afg3l2^M665R/+^* and *Afg3l2^M665R/M665R^* MEFs showed significant reduction in: (i) basal oxygen consumption rate; (ii) maximum respiratory capacity; (iii) spare respiration reserve and (iv) ATP-linked respiration, evaluated using the Seahorse system (Fig. 4A; Supp. Fig. S2A). In addition, *Afg3l2^M665R/+^* MEFs showed reduced extracellular acidification rates, indicating reduced metabolic activity compared to controls (Fig. 4A; Supp. Fig. S2A). The mitochondrial membrane potential (ΔΨm) was significantly decreased in *Afg3l2^M665R/M665R^* MEFs (Fig. 4B) compared to wild-type at steady state, pointing to a defect in the proton pump possibly caused by impaired function of OXPHOS complexes.

**Figure 4.**
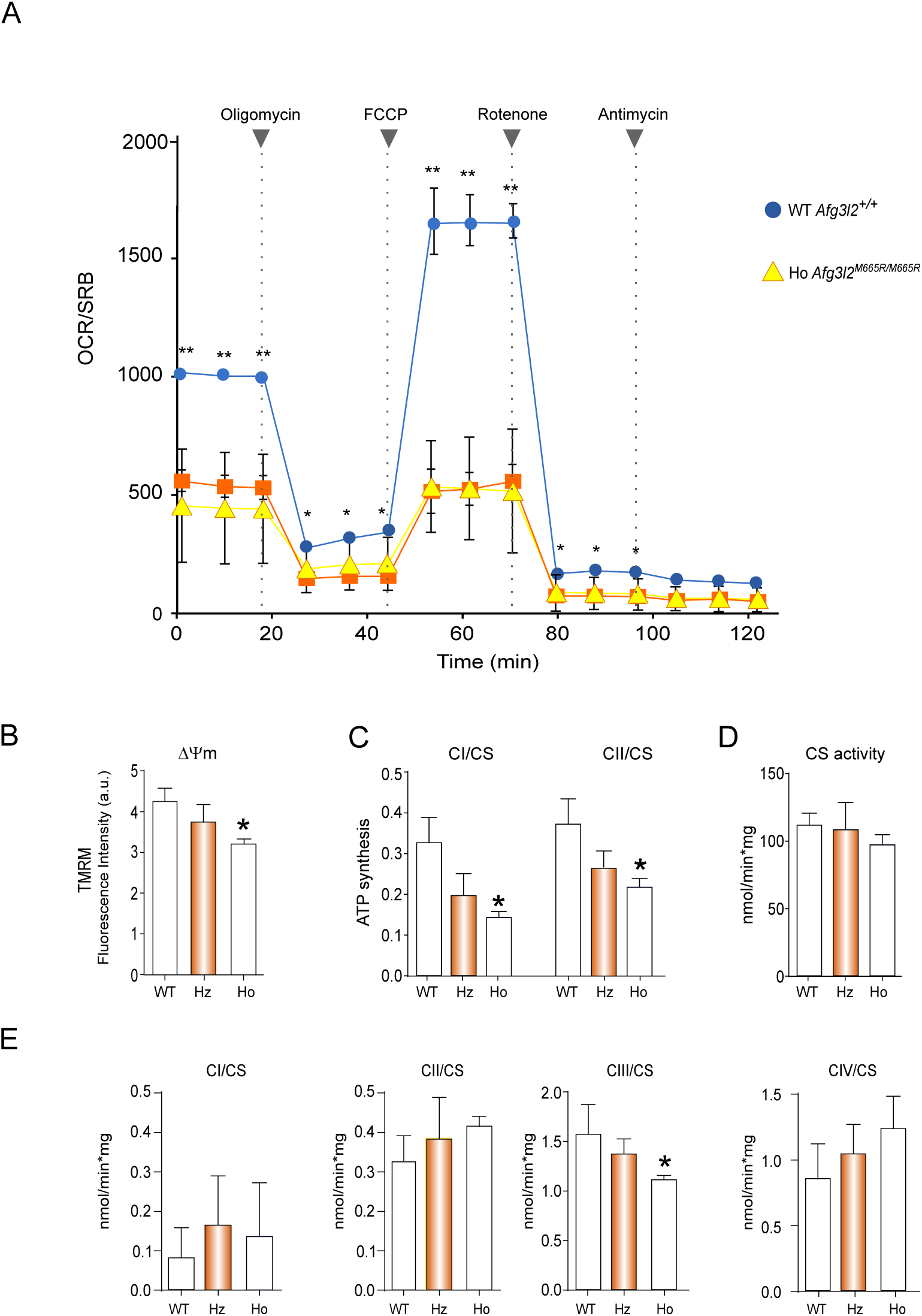
Comparative analysis of mitochondrial bioenergetics in mouse embryonic fibroblasts (MEFs). (A). Oxygen consumption rate (OCR) was measured by Seahorse assay in wild-type (WT), heterozygous (Hz, *Afg3l2^M665R/+^*) and homozygous (Ho, *Afg3l2^M665R/M665R^*) mutant MEFs. Grey dotted lines show the time point and order in which the indicated compounds (Oligomycin → FCCP → Rotenone → Antimycin) were added. (B). Results of TMRM (Tetramethylrhodamine, methyl ester) assay to measure mitochondrial membrane potential in WT, Hz and Ho MEFs (* is *p* < 0.05, Student’s t-test). (C). ATP synthesis assay in digitonin-permeabilized cells comparing CI-driven and CII-driven ATP synthesis in WT, Hz and Ho MEFs (* is *p* < 0.05, Student’s t-test). (D). Analysis of citrate synthase activity in MEFs. (E). Spectrophotometric analysis in isolated mitochondrial fractions showing normal complex I, II and IV activity. Complex III is significantly reduced (30%) in homozygous (Ho) mutants with respect to WT and Hz (* is *p* < 0.05, Student’s t-test).

Reduced ATP-linked respiration was confirmed by ATP synthesis assay performed in digitonin-permeabilized cells. Homozygous mutant MEFs exhibited a significant decrease of Complex I-driven and Complex II-driven ATP synthesis, indicating that the p.Met665Arg Afg3l2 variant impairs respiratory complex chain activity (Fig. 4C). By spectrophotometric analysis of isolated mitochondrial fractions, we found normal levels of activity in Complex I, II and IV in both *Afg3l2^M665R/+^* and *Afg3l2^M665R/M665R^* MEFs. However, there was a 30% reduction of Complex III activity, which was statistically significant, in mitochondria isolated from *Afg3l2^M665R/M665R^* MEFs. Citrate synthase activity was comparable in all three genotypes (Fig. 4D-E), indicating that there were no overall differences in mitochondrial mass.

To evaluate mitochondrial protein maturation together with the assembly and stability of OXPHOS complexes, we analyzed mitochondrial *in vitro* translation, the steady-state levels of selected OXPHOS complex subunits and of fully assembled respiratory complexes by native PAGE. No differences were found between MEFs in terms of *de novo* mitochondrial protein synthesis assessed *in organello* after incorporation of ^35^S-Met/Cys (Supp. Fig. S2B), indicating proper synthesis of mitochondrial proteins in the presence of mutant Afg3l2. The steady-state levels of respiratory complex subunits in MEFs were similar for all genotypes (Supp. Fig. S2C). Fully assembled Complex III was reduced in *Afg3l2^M665R/M665R^* MEFs, possibly explaining the reduced Complex III redox activity found in these cells, whereas the other respiratory complexes were expressed in normal amounts (Supp. Fig. 2D).

### Changes in mitochondrial dynamics induced by the p.Met665Arg mutation

To determine the impact of the p.M665R mutation on mitochondrial morphology, MEFs were treated with MitoRed to visualize fluorescently stained mitochondria. *Afg3l2^M665R/M665R^* MEFs showed fragmented mitochondria, reminiscent of observations made in both *Afg3l2^Emv66/Emv66^* MEFs and mutant mice (Maltecca et al. 2012) whereas *Afg3l2^M665R/+^* MEFs showed intermediate/tubular mitochondria (Fig. 5A, B).

**Figure 5.**
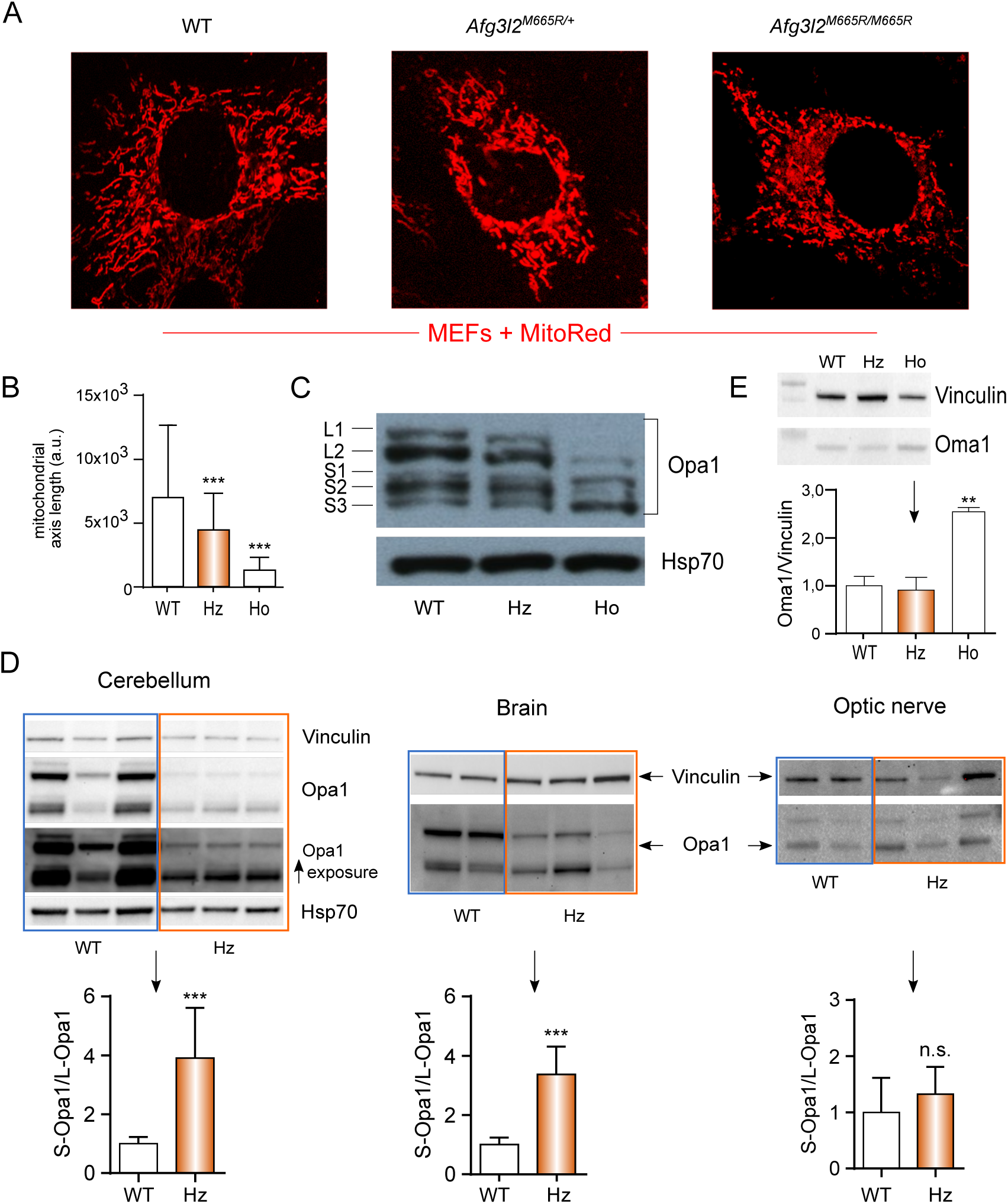
Mitochondrial dynamics in MEFs. (A). Representative images obtained by live confocal microscopy showing mitochondrial morphology in wild-type, *Afg3l2^M665R/+^* (Hz) and *Afg3l2^M665R/M665R^* (Ho) MEFs after staining with mitoRED fluorescent probe. (B). Morphometric analysis of mitochondrial morphology: mitochondria from 50 MEF cells per genotype were measured along the length of the major mitochondrial axis. Histograms represent mean ± SD of four independent experiments (a.u., arbitrary units). WT vs. either Hz or Ho (*** is *p* < 0.0001, Student’s t-test, two-tailed). (C). Immunoblot of MEF total lysates stained for Opa1; Hsp70, loading control. (D). Immunoblot analysis of Opa1 processing in homogenates of neuronal tissues from WT and heterozygous *Afg3l2^M665/+^* mice. Histograms below gels show the ratio between short and long Opa1 isoforms. (E). Immunoblot showing the 60 kDa form of pre-pro-Oma1 (*** is *p* < 0.0001, Student’s t-test, two-tailed). Histogram below the gel shows Oma1/vinculin ration. Vinculin, loading control.

MEF expression of proteins that regulate mitochondrial fission and fusion were analysed by western blot. The GTPase Opa1 (Optic atrophy 1) is a major regulator of mitochondrial morphology: the long isoforms (L-Opa1) promote mitofusion whereas the proteolytically-generated short forms (S-Opa1) instead trigger mitochondrial fragmentation and fission. *Afg3l2^M665R/M665R^* MEFs showed a remarkable decrease in LOpa1, pointing to increased proteolytic processing to S-Opa1 (Fig. 5C). We also analysed expression of Drp1 (pro-fission) and the mitofusins Mfn1 and Mfn2 in MEFS from the three genotypes, but found no major differences in expression (Supp. Fig. S2E).

To return to our mouse model, we next evaluated Opa1 expression in neuronal tissues from wild type and heterozygous *Afg3l2^M665R/+^* mice by western blot. L-Opa1 was greatly decreased in cerebella of *Afg3l2^M665R/+^* mice (Fig. 5D) and in total brain lysate, but not in optic nerves. The S-Opa1/L-Opa1 ratio is shown beneath each gel (Fig. 5D). Overall, these results suggest that mitochondrial fragmentation, specifically in cells of the brain and cerebellum, is altered in our mouse model of SCA28.

### Chloramphenicol treatment rescues the mitochondrial phenotype in *Afg3l2^M665R/M665R^* MEFs

The increase in Opa1 processing with consequent fragmentation of the mitochondrial network, described above in our murine model, is proposed to be a primary event in our model of pathogenesis leading to SCA28. It has been reported that *m-*AAA malfunction triggers activation of OMA1, a back-up protease (ATP-independent) that cleaves OPA1 at the S1 cleavage site, leading to the formation of S1 and S3 isoforms (Anand et al. 2014; Ehses et al. 2009; MacVicar and Langer 2016; Quiros et al. 2012). Accordingly, in our MEF model we found that both S1 and S3 forms of Opa1 were increased in homozygous mutant MEFs by western blotting (Fig. 5C). It is also known that OMA1 is synthesized as a pre-pro-protein of 60 kDa, which accumulates in parallel with the increase in cleaved OPA1 variants (Head et al. 2009). Pre-pro-OMA1 undergoes proteolytic processing after mitochondrial import, generating the stress-sensitive form, pro-OMA1 (40 kDa). This in turn is cleaved to generate the active form of OMA1, which is responsible for OPA1 processing (Baker et al. 2014). Very recently, AFG3L2 has been shown to play a major role in converting pre-pro-OMA1 to pro-OMA1 (Consolato et al. 2018). In line with these observations, we found that in our MEF model, murine pre-pro-Oma1 (60 kDa) was increased in homozygous mutants MEFs compared to WT (Fig. 5E) by western blot.

A consequence of AFG3L2 malfunction is that mitochondrial protein synthesis leads to OMA-1 activation (Arlt et al. 1996; Richter et al. 2015). Similarly, we expected mitochondrial protein synthesis to perturb mitochondrial quality control in our mutant mouse model bearing dysfunctional *m*-AAA complexes containing Afg3l2 p.Met665Arg subunits. Prokaryotic and mitochondrial protein synthesis machineries are similar, so antibiotics such as chloramphenicol (CAM) that target bacterial ribosomes often affect mitochondrial protein synthesis too. We tried blocking mitochondrial protein synthesis in MEFs with CAM: remarkably, 24h CAM treatment completely restored L-Opa1 levels to normal in homozygous mutant MEFs (remaining elevated up to 24 h after treatment ended) and also rescued the fragmented mitochondrial phenotype (Fig. 6A-C). The results obtained with mice and MEFs bearing SCA28-type mutations are consistent with perturbed substrate recognition by Afg3l2, leading to mitochondrial proteotoxicity through gradual accumulation of incorrectly processed or improperly degraded proteins.

**Figure 6.**
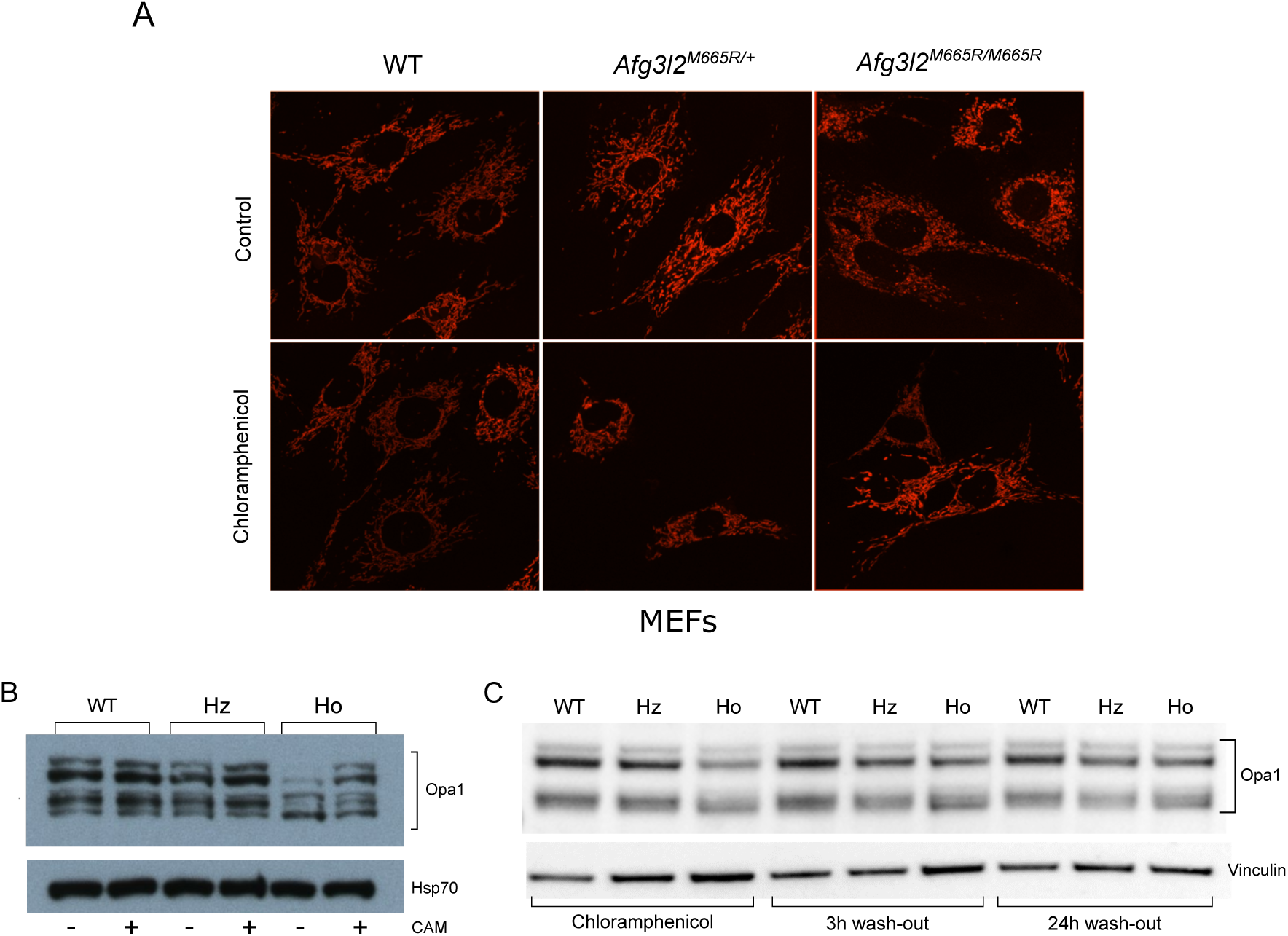
Rescue of mitochondrial dynamics by chloramphenicol treatment. (A). Representative images obtained by live confocal microscopy showing mitochondrial morphology in MEFs stained with mitoRED probe. Cells were untreated or treated with 200 μg/ml chloramphenicol. (B). Immunoblot analysis of total lysates obtained from MEFs untreated or treated with CAM; Hsp70 was used as loading control. (C). Immunoblot analysis of MEF total lysates treated with chloramphenicol, 3h and 24h after CAM wash-out. Vinculin was used as loading control.

## DISCUSSION

Since the discovery of *AFG3L2* as the causative gene of SCA28, understanding the molecular basis of this neurodegenerative disease has been a major goal. The route we selected here was to produce a genetically engineered mouse that mirrored the human disease as closely as possible. At the outset, the idea of modelling a mild human adult-onset ataxia in mice was not without risk, as their short life-span might not allow the time needed for the phenotype to develop.

A number of murine models targeting *Afg3l2* have been developed, with genetic modifications not present in human SCA28. These models vary mechanistically from haploinsufficiency (*Afg3l2^Emv66/+^*) to complete deficiency *in toto* (*Afg3l2^Emv66/Emv66^*, *Afg3l2^par/par^*) to selective deficiency in Purkinje cells (*Afg3l2*^*PC*-KO^). These models have been successful in creating severe disease phenotypes that recapitulate in part the neurological manifestations found in SCA28 (Almajan et al. 2012; Maltecca et al. 2008; Maltecca et al. 2009; Martinelli et al. 2009). However, none of the current models harbor a single-copy patient mutation. Thus the new knock-in mouse model reported here, in which one allele of *Afg3l2* is modified to carry the c.1994T>G:p.Met666Arg point mutation (p.Met665Arg in mouse) derived from a SCA28 patient, can claim to genetically mirror the situation in humans.

How faithfully does the murine phenotype of the *Afg3l2^M665R/+^* mouse recapitulate the human disease? In patients, the ataxic phenotype generally starts in adulthood and progresses very slowly (patients are usually not confined to a wheelchair until into their sixties or seventies). That our knock-in mice did develop a mild form of ataxia at 18 months of age is consistent with the human pattern of disease, just as the late-onset is consistent with the differences in human-mouse chronobiology. The mouse motor phenotype was detected by the beam walking test. This is the most sensitive method to reveal fine coordination defects and motor ataxia due to cerebellar dysfunction. However, the other motor tests were non-discriminatory. One possible interpretation is that although early cerebellar damage may be present, it does not yet affect most motor functions.

How does the *Afg3l2^M665R/+^* mouse compare to other mouse strains with modified *Afg3l2*? The knock-in mouse has a milder phenotype than that of the haploinsufficient strains: (i) in *Afg3l2^Emv66/+^* mice, gait alterations develop at 4 months (Maltecca et al. 2009); (ii) in *Afg3l2^PC-KO^* mice, gait unsteadiness develops at 6-8 weeks (Almajan et al. 2012), and (iii) in *Spg*^*7-/-*^*Afg3l2^Emv66/+^* mice, altered coordination develops at 6 weeks (Martinelli et al. 2009). Thus, in terms of late-onset mild ataxia, the *Afg3l2^M665R/+^* mouse appears to more faithfully represent the SCA28 phenotype.

In SCA28, there is MRI evidence of cerebellar atrophy with Purkinje cell loss in almost all patients analysed. Despite the detailed investigation, we found no evidence of cerebellar pathology in the knock-in mice. On the other hand, this is consistent with the mildness of the motor deficits observed. Purkinje cell numbers were normal, without the dark cell degeneration described in the *Afg3l2^Emv66/+^* mice (Maltecca et al. 2009). We postulate that the most likely cause of motor incoordination in *Afg3l2^M665R/+^* mice must be a qualitative rather that quantitative alteration of Purkinje cells. An indication of altered function was their tendency towards increased action potential firing. This tendency could be linked to impaired buffering of the ionic currents underlying the firing activity, and could represent an initial sign of Purkinje cell dysfunction, leading to their incapacity to depolarize correctly.

Purkinje cell dysfunction points to the crucial role of AFG3L2 in maintaining functional organelles and regulating mitochondrial dynamics (Koppen and Langer 2007; Koppen et al. 2007). With their complex dendritic arborizations and synapses, Purkinje cells require a healthy and extensive mitochondrial network for ATP synthesis, axonal transport, and to buffer calcium influx. This network is maintained by a delicate balance between the opposing forces of mitochondrial fusion and fission. A particular feature of Purkinje cell mitochondria is that they have a longer lifespan than those of other tissues, making them more vulnerable to the cumulative effects of oxidative, proteotoxic, DNA and other forms of damage (Chen et al. 2007).

How does Afg3l2 - p.Met665Arg affect the function of the *m-*AAA protease and lead to mitochondrial dysfunction? We addressed this question using the mouse embryonic fibroblast model, which revealed various mitochondrial alterations. Firstly, there was evident increased mitochondrial fragmentation in homozygous *Afg3l2^M665R/M665R^* MEFs, whereas *Afg3l2^M665R/+^* MEFs displayed an intermediate/tubular phenotype. Fragmentation of the mitochondrial network is known to be caused by an imbalance in proteolytic cleavage of Opa1, which together with the back-up protease Oma1, is one of the determinants of mitochondrial survival (Rugarli and Langer 2012). Our findings of increased fragmentation were corroborated by biochemical evidence of overproduction of the short isoform of Opa1 (in particular S1 and S3 isoforms) in *Afg3l2^M665R/M665R^* and *Afg3l2^M665R/+^* MEFs. Extensive mitochondrial fragmentation was also detected in *Afg3l2^Emv66/Emv66^* MEFs (Maltecca et al. 2009). Proceeding from the MEF cellular model to the whole organism, we also observed impaired Opa1 processing in cerebellar lysates of 18-month old *Afg3l2^M665R/+^* mice, suggesting this could be one of the early events in SCA28 pathogenesis, although patient mitochondria have yet to be analysed. Increased processing of Opa1 could also play an important role in the perinatal lethality of the homozygous knock-in mice. Opa1 is central for cardiac function, especially during cardiomyocyte differentiation and heart development (Wai et al. 2015). Like Opa1, Afg3l2 is also highly expressed in the developing heart (http://www.genecards.org/cgibin/carddisp.pl?gene=AFG3L2&keywords=afg3l2), possibly linking the two proteases in the perinatal lethality of the knock-in mouse, which will be a future avenue of research.

Altered Opa1 cleavage has also been associated with enhanced apoptosis (Jiang et al. 2014; Rugarli and Langer 2012), altered morphology of mitochondrial *cristae*, and decreased cell respiration and ATP production (Anand et al. 2014). This is consistent with the impaired bioenergetics observed in *Afg3l2^M665R/M665R^* MEF mitochondria that revealed significantly reduced respiration and ATP synthesis. Altered Opa1 processing in *Afg3l2^M665R/M665R^* MEFs may be linked to the decrease in mitochondrial membrane potential compared to WT MEFs, which has been shown to induce activation of the ATP-independent zinc metalloprotease Oma1 which, in turn, promotes Opa1 proteolytic cleavage (Zhang et al. 2014). Indeed, increased pre-pro-Oma1 (60 kDa form) was detected in *Afg3l2^M665R/M665R^* MEFs, supporting its activated state (Head et al. 2009; Korwitz et al. 2016; Consolato et al. 2018).

Based on the results presented here, we hypothesize that Afg3l2 p.Met665Arg reduces the processing capability/rate of *m-*AAA, resulting in failure to turnover damaged or misfolded mitochondrial proteins, together with the accumulation of *de novo* synthesized proteins *in organello*. The burden of newly synthesized mitochondrial proteins may provoke inner membrane depolarization which, in association with protetoxic stress, activates the Oma1 protease. Consequently, Opa1 processing is impaired, fragmentation is enhanced and the mitochondrial network is disrupted, causing a functional alteration of the cell, and ultimately impairing neuronal functions (Fig. 7). Therefore, we decided to block mitochondrial *de novo* protein synthesis using the antibiotic chloramphenicol, a broad spectrum antibiotic that inhibits translation in both mitochondria and prokaryotes (Bulkley et al. 2010; Richter et al. 2015). By blocking mitochondrial protein synthesis, we were able to restore the correct balance between Opa1 isoforms and re-establish the mitochondrial network in *Afg3l2^M665R/M665R^* MEFs. This result demonstrates that the p.Met665Arg mutation in Afg3l2 affects the chaperone and proteolytic functions of the *m-*AAA complex, reducing protein quality surveillance in mitochondria. In agreement with this, bioinformatics analysis of AFG3L2 p.Met666Arg missense change showed that the electrostatic potential difference of the homohexamer complex central pore is altered with respect to wild-type AFG3L2, pointing to a reduction in the capacity of *m*-AAA to deliver substrates to the proteolytic chamber, or to a weaker interaction among monomers (Cagnoli et al. 2010). Chloramphenicol may also be affecting crosstalk between mitochondria and nucleus, as stalled translation might activate a retrograde signal to the nucleus, resulting in a compensatory pathway that inhibits Opa1 cleavage.

**Figure 7.**
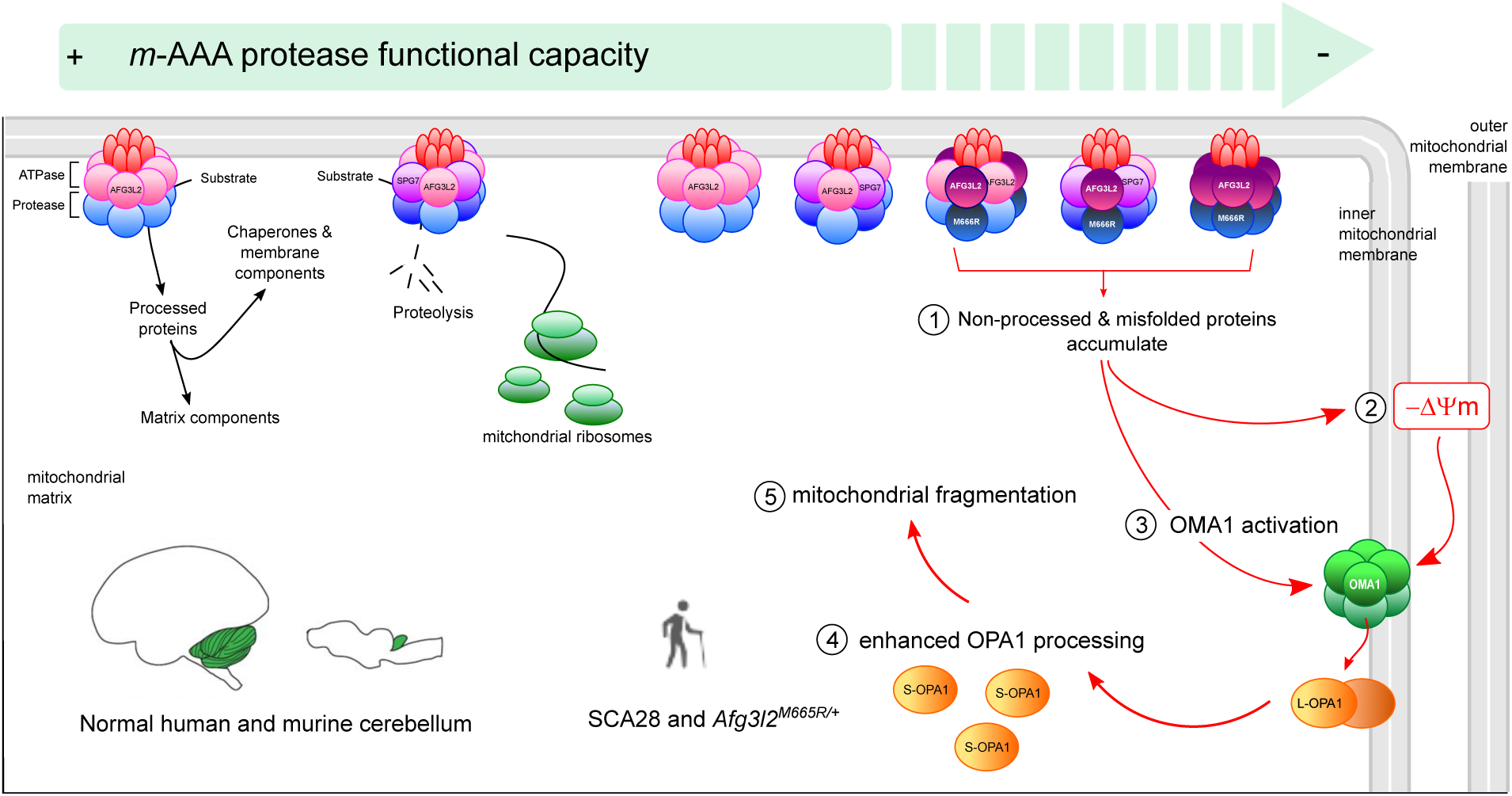
Pathogenic mechanisms triggering SCA28. Mitochondria have an elaborate proteolytic system, present within different subcompartments of the organelle, which coordinates the maturation and processing of newly synthesized proteins and the elimination of non-assembled polypeptides to avoid potential deleterious effects on organelle function. In physiological conditions (left), the *m-*AAA exerts crucial functions in mt protein quality control: (i) it participates in unfolded mt-protein degradation; (ii) it is involved in processing and maturation of mt-proteins; (iii) it has chaperoning activity, helping mt-proteins insert into the inner mitochondrial membrane. The processing capability/rate of the *m-*AAA is reduced or blocked, proportionally to the number of Afg3l2 p.Met665Arg integrated in the complex (right), resulting in (1) the accumulation of non-processed and misfolded proteins in the organelles that lack processing or degradation by the *m-*AAA protease. This burden of newly synthesized mt-proteins derived from impairment of the mitochondrial proteolytic system, activates OMA1 protease (3), directly or through an inner membrane depolarization (2). Consequently, Opa1 processing is enhanced (4), leading to fragmentation of the mitochondrial network (5). Chloramphenicol, acting by inhibiting *de novo* translation in mt-ribosomes, prevents the accumulation of newly synthesized proteins and reduces stress signals in the inner membrane, thus rescuing the cascade of events leading to Opa1 enhanced processing and mt-fragmentation.

Experimental data involving the *m-*AAA protease must take into account that its roles are only partially understood. In mice, Afg3l1 (a pseudogene in humans), a third paralogue of Afg3l2 and Spg7, can modulate *m*-AAA function. Thus, at least five different types of *m-*AAA complexes are formed (Afg3l2 x 6; Afg3l1 x 6; Afg3l2 + Spg7; Afg3l2 + Afg3l1 and Afg3l1 + Spg7) (Koppen et al. 2007) and different hexamers may have distinct substrate or tissue specificities. For example, we know that Afg3l1 is the least expressed of the subunits in all tissues analysed, especially in cerebellum and brain (Sacco et al. 2010). In the presence of a mutated Afg3l2 with impaired protease activity, paraplegin and Afg3l1 subunits could be sequestered in non-functional hetero-hexameric complexes, reducing the amount of functional *m*-AAA complexes. In contrast, *Afg3l2* deficiency would lead to the complete absence of protein, allowing Afg3l1 homo-hexamers and Afg3l1/Spg7 hetero-hexamers to form functional *m-*AAA complexes. Such a compensatory effect that has been demonstrated in *Afg3l2^PC-KO^* mouse oligodendrocytes (Almajan et al. 2012).

In *Afg3l2^M665R/+^* mice, Afg3l2 synthesis is not impaired and a physiological number of *m*-AAA complexes should be formed, with wild-type and mutated subunits participating equally. We propose that a certain quantity of mutant Afg3l2 subunits can be tolerated, allowing normal *m-*AAA function to be maintained (up to a point). Thus we speculate that the heterozygous knock-in mice form a greater number of functional (or partially functional) *m*-AAA hexamers than in the heterozygous knockout mice, explaining the less severe phenotype. Indeed, all the previous mouse models strongly support a gene dosage effect of the three subunits (Afg3l2, Afg3l1 and Spg7) in determining the functional outcome of *m-*AAA hexamers (Ferreirinha et al. 2004; Maltecca et al. 2008; Maltecca et al. 2009; Martinelli et al. 2009; Wang et al. 2016). However, given that we ignore the *m-*AAA subunit stoichiometry and the reciprocal disposition of the subunits (alternate/continuous) in the complex, it remains a speculative hypothesis that warrants further experiment.

The role of Afg3l2 in Spg7 processing may also shed light on the effect of p.Met665Arg. Spg7 must be processed at two cleavage sites to activate its protease activity, the latter being dependent on Afg3l2/Afg3l1 (Koppen et al. 2009). By electron microscopy, sciatic nerves of *Afg3l2^M665R/+^* mice show swollen mitochondria with disrupted *cristae*, corroborating the idea that the Afg3l2-paraplegin complex, highly represented in the peripheral nervous system, is also affected in our knock-in model (Koppen et al. 2007; Sacco et al. 2010).

In conclusion, we have investigated the molecular mechanisms leading to SCA28 with a newly developed knock-in mouse model which we believe better mimicks the human disease than previous models. We propose that mitochondrial proteostasis, leading to slow and toxic accumulation, is the pathogenic triggering event. Our findings open a novel therapeutic route, based on the modulation of mitochondrial translation, to be investigated in future studies.

## MATERIALS AND METHODS

### Preparation of *Afg3l2^M665R/+^* knock-in mouse model

To generate our knock-in mouse model, we used a gene targeting approach to introduce the M665R mutation into *Afg3l2*. The targeting vector was constructed using the recombineering strategy (www.recombineering.ncifcrf.gov) (Copeland et al. 2001) and consisted of a 14.1 kb *NotI-XhoI* fragment from the BAC bmp360E12 (mouse strain 129) containing exon 16 and the flanking intron sequences (5,261 bp upstream and 7,128 bp downstream) of murine *Afg3l2*. In a second vector, we cloned a 0.6 kb region flanking exon 16 into which we introduced, through *SalI-NotI* restriction cloning, the *neo^r^* cassette flanked by two Flippase recognition target sequences. The c.1994T>G point mutation (p.Met665Arg) was introduced by site-directed mutagenesis (Stratagene) together with an *NcoI* restriction site at the 5’ end of exon 16 to allow discrimination between wild-type and mutated alleles by Southern blot. We used recombineering to generate the final vector for electroporation into mouse 129 ES cells. After positive selection with neomycin, 230 clones were harvested; 10 were positive by Southern blot using *NcoI/SmiI*-digested DNA. The Southern blot probe was a ∼300-bp fragment of *Afg3l2* cloned into pUC19 after PCR amplification of mouse genomic DNA (primers 5’-ttcaggtctttcactggcatggt; 5’-gcaacacagtactatacacctccttacg). The probed yields a 17.1 kb band in wild-type DNA, and 13.4 kb in mutant cells. The *neo^r^* cassette copy number was evaluated by real-time PCR (Mancini et al. 2011). Only 2/10 ES cell lines showed a single *neo^r^* insertion and were injected into C57BL/6 blastocysts. The resulting chimeras were then mated to C57BL/6 females to obtain germline transmission. A diagnostic restriction enzyme digest was set up to verify the heterozygous presence of the mutation in mice: exon 16 was amplified using primers mAfg3l2_16F (5’-gctggtgcggtttgcccag) and mAfg3l2_16R (5’-cagcaggtagacactagctaagcaacc). Amplicons were digested with *Hsp92II* that cuts only the wild-type allele. The genotype of mice was confirmed by PCR, followed by Sanger sequencing. The *neo^r^* cassette was eliminated by crossing *Afg3l2^M665R-neo/+^* mice with a mouse strain containing the Flippase recombination enzyme. *Afg3l2^M665R/+^* animals were backcrossed with C57BL/6J for at least seven generations to purify the genetic background. Studies were performed in accordance with the guidelines of the Neuroscience Institute Cavalieri Ottolenghi (NICO) (Orbassano, Italy), after obtaining approval by the Ethics Committee of the University of Torino, and following authorization from the Italian Ministry of Health (Authorization no.: 58/2016-PR).

### Assessment of motor phenotype

Motor tests were performed every two months to detect the onset of neurological signs, starting when mice were 4 months old. The methods were as follows for 18 month old mice. The 18-month-old wild-type and *Afg3l2^M665R/+^* littermates were of similar weight (wild-type: 25.38 ± 1.11 g, n = 11; *Afg3l2^M665R/+^*: 25.98 ± 0.62 g, n =18; *P* > 0.05). For the beam test (Goldowitz 1992), *Afg3l2^M665R/+^* mice (n = 16) and their control littermates (n = 14) were placed on a metal beam (80 cm long x 1 cm wide) suspended above an open cage. Mice were allowed to traverse the beam for four days before the test (habituation). During the test, lateral slips were counted in three complete and consecutive crossings per day, on two consecutive days. The mean number of slips per 80 cm travelled was calculated. Footprint analysis was carried out using wild-type (n = 8) and *Afg3l2^M665R/+^*(n = 8) mice, as described in Hoxha *et al.* (Hoxha et al. 2013). Briefly, a cardboard tunnel was constructed on a transparent plexiglass platform (4 cm wide x 40 cm long). Video footage of mouse walks was obtained and analyzed using ImageJ software. In the accelerating rotarod test, wild-type (n=13) and *Afg3l2^M665R/+^*(n=13) mice were tested on five consecutive days, and on day 10. Each day, after 1 min training at constant speed (4 rpm), mice had three test sessions in which the rod (Mouse Rota-Rod, Ugo Basile Biological Research Apparatus, Comerio, Italy) accelerated continuously from 4 to 65 rpm over 300 s. The time to falling off the rod was recorded.

The optokinetic reflex (OKR) was measured to assess visual function. The OKR was evoked using a rotating optokinetic drum with an 8° vertical grating. After placing the mouse on a central platform in the drum, and allowing 1 min for acclimation, the drum was rotated counterclockwise for 1 min, stopped for 10 s and then rotated clockwise for another minute. Rotation velocity was 10°/s. The mouse was video-recorded and visual head tracking was measured offline (Thaung et al. 2002).

### Histological analyses

Eighteen-month-old *Afg3l2^M665R/+^* (n = 3) and wild-type (n = 3) mice were anesthetized by intraperitoneal injection with ketamine (100 mg/kg body weight) and xylazine (10 mg/kg body weight), and perfusion-fixed with 4% paraformaldehyde in 0.12M phosphate buffer (PB). Brains were removed and immersed in the same fixative at 4°C for 18h and then cryoprotected in 30% sucrose in 0.12M PB. Cerebella were frozen and serially cut on a freezing microtome into 30 μm-thick sagittal sections. To label Purkinje cells, one set of sections was incubated overnight at 4°C with anti-calbindin-D28k polyclonal antibody (CB-38a, 1:3000; Swant, Bellinzona, CH) diluted in PBS with 0.25% Triton X-100 and 1.5% normal goat serum. Immunohistochemical reactions were stained with avidin-biotin-peroxidase (Vectastain ABC Elite kit; Vector Laboratories, Burlingame, CA, USA) and revealed using 3,3′-diaminobenzidine (0.03% in Tris–HCl) as chromogen. A second set of sections was Nissl-stained to gauge the thickness of the molecular and granule cell cerebellar layers, and to quantify neuronal density (cell number/mm^2^) in the deep cerebellar nuclei (DCN). Morphometric analysis of the cerebellar sections was performed by means of the Neurolucida system (MicroBrightField, Williston, VT, USA) and ImageJ software (http://rsbweb.nih.gov/ij/index.html).

### Purkinje cell firing

Cerebellar slices were prepared as previously described (Llinas and Sugimori, 1980 and Edwards *et al*., 1989). Mice were anaesthetized with isoflurane (Isoflurane-Vet, Merial, Italy) and decapitated. The cerebellar vermis was removed and transferred to ice-cold artificial cerebrospinal fluid (ACSF) containing (in mM); 125 NaCl, 2.5 KCl, 2 CaCl_2_, 1 MgCl_2_, 1.25 NaH_2_PO_4_, 26 NaHCO_3_, 20 glucose, bubbled with 95% O_2_/5% CO_2_ (pH 7.4). Parasagittal cerebellar slices (200 *μ*m thickness) were obtained using a vibratome (Leica Microsystems GmbH, Wetzlar, Germany) and kept for 1 h at 35°C and then at 31°C. Single slices were placed in the recording chamber, which was perfused at a rate of 2–3 ml/min with ACSF bubbled with the 95% O_2_/5% CO_2_. All recordings were performed at 31±1°C temperature. For cell-attached recording, the pipette was filled with 0.9 % NaCl, 0.2 *μ*m filtered. Recordings were made using an EPC‐10 patch‐clamp amplifier (HEKA Elektronik, Lambrecht/Pfalz, Germany). Gabazine (SR 95531, 20 μM), DAP5 (50 μM) and NBQX (10 μM) were added to the perfusate to inhibit the GABA_A_ and ionotropic glutamate receptors of Purkinje cells, respectively. Data was derived from the analysis of 3 - 5 animals. All drugs were purchased from Abcam.

### Electron microscopy

#### Sciatic nerve

To prepare sciatic nerve sections, 18-month-old wild-type mice (n= 5) and *Afg3l2^M665R/+^*(n= 5) were anaesthetized by intraperitoneal injection with ketamine (100 mg/kg body weight) and xylazine (10 mg/kg body weight). Using an operative microscope, the sciatic nerve was carefully exposed and removed. Nerves were immediately fixed by immersion in 2.5% glutaraldehyde in 0.1 M phosphate buffer (pH 7.4) for 4 - 6 h at 4°C. Samples were postfixed in 2% osmium tetroxide for 2 h and dehydrated in ethanol from 30% to 100%. After two passages of 7 min each in propylene oxide and overnight in a 1: 1 mixture of propylene oxide and Glauerts’ mixture of resins, sciatic nerves were embedded in equal parts of Araldite M and the Araldite Harter resin, with further addition of 0.5% dibutylphthalate plasticizer. Finally, 2% of accelerator 964 was added to the resin in order to promote the polymerization of the embedding mixture.

#### Light microscopy and design-based quantitative morphology of sciatic nerves

From each nerve, 2.5-μm-thick series of semi-thin transverse sections were cut using an Ultracut UCT ultramicrotome (Leica Microsystems, Wetzlar, Germany) and stained with 1% Toluidine blue for high resolution light microscopic examination and design-based stereology. A DM4000B microscope equipped using a DFC320 digital camera and an IM50 image manager system (Leica Microsystems) was used for stereology of nerve fibers. On one randomly selected toluidine blue stained semi-thin section, the total cross-sectional area of the whole nerve was measured by light microscope and 12 - 16 sampling fields were selected by systematic random sampling (Geuna 2000; Geuna et al. 2000; Larsen 1998). In each sampling field, a two-dimensional dissector procedure, which is based on sampling the “tops” of fibers, was adopted in order to avoid the “edge effect” (Geuna 2000). Using this procedure, the total number of myelinated fibers (N) was estimated. In addition, both fiber and axon areas were measured, and the diameter of fiber (D) and axon (d) were calculated. These data were used to calculate myelin thickness [(D−d)/2], myelin thickness/axon diameter ratio [(D−d)/2d], and axon/fiber diameter.

#### Quantitative analysis of mitochondria in sciatic nerves

Ultra-thin (70 nm) sections were cut from the sciatic nerves using the same ultramicrotome described for the high resolution light microscopy (above). Nerve sections were stained with saturated aqueous solution of uranyl acetate and lead citrate. Ultra-thin sections were analyzed using a JEM-1010 transmission electron microscope (JEOL, Tokyo, Japan) equipped with a Mega-View-III digital camera and a Soft-Imaging-System (SIS, Münster, Germany) for computerized acquisition of the images. On one randomly selected ultra-thin section, 10 - 15 fields were selected using a systematic random sampling protocol, with a magnification of ×15000. In each sampling field, the number of impaired and unimpaired mitochondria was estimated as %, based on their morphological features such as the shape of mitochondria, the morphology of the *cristae* and evidence of swelling.

#### Cerebellar and lumbar spinal cord electron microscopy

After excision of sciatic nerves, animals were perfused-fixed with 3% glutaraldehyde diluted in 0.1 M Sorensen phosphate buffer (pH 7.4) to prepare cerebellar and lumbar spinal cord sections. Bioptic fragments obtained by cerebellum and lumbar spinal cord were immersed in 3% glutaraldehyde diluted in 0.1 M Sorensen phosphate buffer for 4 hours at 4°C, washed x3 for 30 min in Sorensen phosphate buffer, and post-fixed in 1% Osmium Tetroxide. After dehydration through an ascending series of ethanol solutions, samples were embedded in Durcupan (Durcupan, Fluka, Milan, Italy). Ultra-thin sections were obtained with an Ultracut ultramicrotome (Reichert Ultracut R-Ultramicrotome, Leika, Wien, Austria) and stained with uranyl acetate/lead citrate before observation by a Jeol CX100 transmission electron microscope (Jeol, Tokyo, Japan).

### Mouse embryonic fibroblasts (MEF) production and analysis

#### Generation of MEFs

Primary MEFs were established from embryos obtained on day E13.5 from wild-type, KI-hz *Afg3l2^M665R/+^* and KI-ho *Afg3l2^M665R/M665R^* mice using a standard protocol (Sharpless 2006) with immortalization by SV40 plasmid transfection followed by 300 µg/ml G418 antibiotic selection (Schuermann 1990).

#### Analysis of mitochondrial morphology

Mitochondrial network morphology was analysed using the mito-RED probe (Invitrogen) and live imaging confocal microscopy. An average of 150 cells was analyzed for each experimental condition. Mitochondrial length was measured using ImageJ software tools. Experiments were repeated at least three times, and two operators independently analysed the images. The t-student test on raw data was to calculate significance (*p* < 0.05).

#### Western blot analysis

Total cell or tissue lysates were prepared by protein extraction in RIPA buffer. Twenty μg of proteins were separated by electrophoresis on 4-12% Bis-Tris Protein Gels (Life Technologies). Anti-Afg3l2 (14631-1AP, polyclonal rabbit) and anti-Oma1 (17116-1AP, polyclonal rabbit, specific for pre-pro-Oma1 of 60 kDa) antibodies were purchased from Proteintech. Anti-Vinculin (Millipore, #AB6039) or anti-β-actin (Lifespan, LS-C51570) were used as loading controls. For Opa1 band processing, we performed standard western blotting with total cell or tissue lysates prepared by protein extraction using a Triton-containing buffer (Maltecca et al. 2012). Twenty μg (MEFs) and 5 μg (tissues) were separated by electrophoresis on 8% Bolt™ Bis-Tris Plus Gels (Life Technologies). Anti-Opa1 antibody was purchased from BD Transduction Laboratories #612606 (Franklin Lakes, NJ, USA). Anti-Viculin was used as loading control.

### Analysis of mitochondrial bioenergetics

#### Preparation of crude mitochondria

Dissected tissues were promptly chopped and homogenized (30 strokes) in 10 volumes/g wet tissue of mannitol/sucrose buffer (220 mM D-mannitol, 70 mM sucrose, 20 mM HEPES, 1 mM EDTA and 0.1% BSA, pH 7.2) in a glass-Teflon pestle. The crude homogenate was centrifuged at 500 g for 10 min at 4°C. The low-speed supernatant was centrifuged at 10,000 g for 10 min at 4°C. The resulting mitochondrial pellet was carefully re-suspended to about 10–20 mg/ml in mannitol/sucrose buffer. Protein concentration was determined by Bradford’s method (BioRad, Hercules, CA, USA) with BSA as the standard.

#### Respiratory complex activity

Mitochondrial respiratory chain complex activity was assayed and normalized for protein content and citrate synthase (CS) activity as previously described (Ghelli et al. 2013). All drugs used for biochemical analyses were from Sigma-Aldrich (St Louis, MO, USA).

#### Native gel electrophoresis

To analyze isolated native complexes, mitoplasts were suspended in a buffer containing 750mM aminocaproic acid, 50 mM Bis-Tris, pH 7.0, and solubilized with 20% n-Dodecyl beta-D-maltoside (DDM), corresponding to a DDM/protein ratio of 2.5 (g/g). The suspension was incubated on ice for 10 min and then centrifuged at 13,000 g for 15 min (Ghelli et al. 2013). Solubilized mitochondrial proteins (80 µg) were separated on native 4 - 16% Bis-Tris Gel (Invitrogen), as detailed before (Ghelli et al. 2013). Gels were run as high resolution Clear Native- and Blue Native-PAGE (hrCN- and BN-PAGE). After electrophoresis, samples in CN-PAGE gels were processed for CI in-gel-activity (CI-IGA); samples in BN-PAGE gels were used for western blotting. Immunodetection was carried out by incubating blots with antibodies directed against the NDUFA9 subunit of CI (Thermo Fisher Scientific), SDHA subunit of CII (Thermo Fisher Scientific), UQCRC2 subunit of CIII (AbCAm, Cambridge, UK), and COXIV of CIV (AbCam, Cambridge, UK).

#### Mitochondrial Respiration assay

Oxygen consumption rate (OCR) in adherent cells was measured with by XFe24 Extracellular Flux Analyzer (Seahorse Bioscience, Billerica MA, USA), as previously described (Iommarini et al. 2014). Briefly, cells were seeded in XFe24 cell culture microplates at 2×10^4^ cells/well in 200 μl complete medium and incubated at 37°C in 5% CO_2_ for 24 h. Assays were initiated by replacing the growth medium in each well with 670 μl of unbuffered DMEM-high glucose supplemented with 1 mM Napyruvate pre-warmed at 37°C. Cells were incubated at 37°C for 30 min to allow temperature and pH to equilibrate. After obtaining the OCR baseline measurement, 1μg/ml oligomycin, 0.25 μM FCCP ((2- [2-[4-(trifluoromethoxy)phenyl]hydrazinylidene]-propanedinitrile), 1μM rotenone and 1μM antimycin A were sequentially injected. At the end of each experiment, the medium was removed and protein content was determined by sulforhodamine B (SRB) assay. OCR data (pmol/min) were normalized to SRB absorbance (Iommarini et al. 2014). First, the basal oxygen consumption rate (basal respiration) was measured. Oligomycin inhibited ATP synthase activity, which led to the development of a proton gradient that inhibited electron flux and revealed the state of the coupling efficiency. FCCP uncoupled the respiratory chain and revealed the maximal capacity for reducing oxygen. The spare respiratory capacity was calculated by subtracting the basal respiration from the maximal respiration. Finally, rotenone combined with antimycin A was injected to inhibit the flux of electrons through complexes I and III; the remaining oxygen consumption rate was primarily due to non-mitochondrial respiration.

### Mitochondrial ATP synthesis assay

The rate of mitochondrial ATP synthesis was measured in digitonin-permeabilized cells as previously described (Ghelli et al. 2013). Briefly, after trypsinization, cells (10×10^6^/ml) were suspended in buffer (150 mM KCl, 25 mM Tris–HCl, 2 mM EDTA, 0.1% BSA, 10 mM potassium phosphate, 0.1 mM MgCl_2_, pH 7.4), maintained at room temperature for 15 min, then incubated with 50 μg/ml digitonin until 90-100% of cells were positive by Trypan blue staining. Aliquots (3×10^5^) of permeabilized cells were incubated in the same buffer as above in the presence of the adenylate kinase inhibitor P^1^,P^5^-di(adenosine-5′) pentaphosphate (0.1 mM) and CI substrates (1 mM malate plus 1 mM pyruvate), CII substrate (5 mM succinate plus 1 μM rotenone) or CIII substrate (50 µM DBH_2_ plus 1 μM rotenone and 5 mM malonate). After the addition of 0.2 mM ADP, chemiluminescence was determined as a function of time with a luminometer (Sirius-L, Berthold, Germany). The chemiluminescence signal was calibrated with an internal ATP standard after the addition of 1 μM oligomycin. The rate of ATP synthesis was normalized to protein content and CS activity.

### Mitochondrial Δψ_m_ assay

Mitochondrial Δψ_m_ was assessed by FACS analysis, after incubating cells with 20nM tetra-methylrhodamine methyl ester perchlorate (TMRM) (Molecular Probes) for 30 min at 37°C. After this, mitochondria were fully depolarized by addition of 5 µM CCCP, as internal control. Data were analysed by WinList software (Verity Software House).

### Mitochondrial in vitro translation

Mitochondrial protein synthesis was analysed in cultured cells by metabolic labelling with [^35^S]methionine/cysteine in the presence of anisomycin to inhibit cytoplasmic ribosomes, as described in (Richter et al. 2015).

## SUPPORTING INFORMATION LEGENDS

**Figure S1. Comparative analysis of whole embryos and selected tissue histology of E16.5 embryos** (A). Photograph showing intact E16.5 embryos isolated from homozygous mutant, wild-type and heterozygous mutant lineages. (B, C). Histological section of heart derived from E16.5 WT (left panel) and homozygous mutant (right panel) mice. (D, E). Cerebellar sections from WT (left panel) and homozygous mutant mice (right panel). Cerebellum is outlined in red dotted line. Hematoxylin/eosin stain.

**Figure S2. Seahorse assay for mitochondrial function and mitochondrial protein analysis in MEFs** (A). Histograms represent results of seahorse assays in wild-type (WT), heterozygous *Afg3l2^M665R/+^* (Hz) and homozygous *Afg3l2^M665R/M665R^* embryonic fibroblasts. Double asterisks (**) shows statistically significant reduction in basal and maximal oxygen consumption rate (OCR), spare OCR capacity and ATP-linked OCR in both Hz and Ho mutants with respect to WT. Single asterisk (*) shows proton leakage reduction was statistically significant only in Hz compared to WT (*p* < 0.05, Student’s t-test, two tailed). (B). Analysis of mitochondrial protein synthesis by pulsed ^35^S metabolic labelling in WT and Hz MEFs (left panel), and WT and Ho MEFs (right panel). Labelled proteins were separated by 12-20% SDS-PAGE and visualized by autoradiography. To control for protein loading, part of the gels was stained with Coomassie blue. (C). Immunoblot analysis of mitochondrial lysates from MEFs showing the steady-state levels of respiratory complex subunits. β-actin, loading control. (D). In-gel activity (IGA) of complex I and blue native (BN) page immunoblot analysis of fully assembled respiratory complexes II, III and IV. A mild reduction in CIII assembly was observed in Ho compared to WT (asterisk*). Mitochondrial proteins Sdha and Vdac were use as loading controls. (E). Immunoblot analysis showing the steady-state levels of fission/fusion regulatory proteins Mitofusin 1 and 2 (Mfn1, 2), and Dynamin-related protein 1 (Drp1) in MEF total lysates.

